# CodonBERT: Large Language Models for mRNA design and optimization

**DOI:** 10.1101/2023.09.09.556981

**Authors:** Sizhen Li, Saeed Moayedpour, Ruijiang Li, Michael Bailey, Saleh Riahi, Lorenzo Kogler-Anele, Milad Miladi, Jacob Miner, Dinghai Zheng, Jun Wang, Akshay Balsubramani, Khang Tran, Minnie Zacharia, Monica Wu, Xiaobo Gu, Ryan Clinton, Carla Asquith, Joseph Skaleski, Lianne Boeglin, Sudha Chivukula, Anusha Dias, Fernando Ulloa Montoya, Vikram Agarwal, Ziv Bar-Joseph, Sven Jager

## Abstract

mRNA based vaccines and therapeutics are gaining popularity and usage across a wide range of conditions. One of the critical issues when designing such mRNAs is sequence optimization. Even small proteins or peptides can be encoded by an enormously large number of mRNAs. The actual mRNA sequence can have a large impact on several properties including expression, stability, immunogenicity, and more. To enable the selection of an optimal sequence, we developed CodonBERT, a large language model (LLM) for mRNAs. Unlike prior models, CodonBERT uses codons as inputs which enables it to learn better representations. CodonBERT was trained using more than 10 million mRNA sequences from a diverse set of organisms. The resulting model captures important biological concepts. CodonBERT can also be extended to perform prediction tasks for various mRNA properties. CodonBERT outperforms previous mRNA prediction methods including on a new flu vaccine dataset.

## 1 Introduction

mRNA vaccines have emerged as a high potency, fast production, low-cost, and safe alternative to traditional vaccines [1, 2, 3, 4]. mRNA vaccines are currently being developed for a broad range of human viruses and bacteria including SARS-CoV-2, influenza, Zika, Chlamydia, and more [5, 6, 7, 8]. They are also being investigated as potential treatments for several diseases including lung cancer, breast cancer, and melanoma [9, 10].

The expression level of a vaccine directly affects its potency, ultimate immunogenicity, and efficacy [11]. The higher the level of expression of the antigenic protein encoded by the mRNA sequence, the smaller amount of the vaccine is needed to achieve the desired immune response, which can make the vaccine more cost-effective and easier to manufacture [1]. Consequently, using a lower dose can help reduce reactogenicity [12] and maintain the immune response over a longer period [13], leading to better safety and efficacy.

A human protein with an average length of 500 amino acids can be encoded by roughly 3^500^ different codon sequences. While only one of those is encoded in the virus or DNA of interest, this is not necessarily the optimal sequence for a vaccine. The classical method to find the optimal mRNA sequence is codon optimization, which selects the most optimal codon for each amino acid using the codon bias in the host organism [14]. This method has been widely applied including for optimizing recombinant protein drugs, nucleic acid therapies, gene therapy, mRNA therapy, and DNA/RNA vaccines [15, 16, 17]. However, codon optimization alone does not consider several key properties that impact protein expression [18]. For instance, RNA structural properties (e.g., stem loops and pseudoknots) have been shown to play a major role for non-coding RNAs (such as riboswitches or aptamers) [19, 20].

While mRNA sequence heavily influences cellular RNA stability [21, 22], secondary structure can also impact mRNA stability in solution and modulate protein expression [13, 23, 24, 25]. For example, replacing a codon with a synonymous codon can alter the local base-pairing interactions and affect nearby structural motifs [26, 27]. Thus, optimizing each codon independently is not sufficient to generate highly expressed proteins.

Pre-training a large language model (LLM) based on large-scale unlabeled text, followed by fine-tuning, has been widely adopted for natural language processing [28, 29, 30, 31]. Recently, this concept has been scaled to biological sequences (protein, DNA, and RNA) [32, 33, 34, 35, 36]. Such models can be used to embed nucleotides and use these embeddings for downstream supervised learning tasks. However, as we show, such LLMs may not be ideal for predicting protein expression due to their focus on individual nucleotides and non-coding regions.

To address these limitations, we developed CodonBERT, an LLM which extends the BERT model and applies it to the language of mRNAs. CodonBERT uses a multi-head attention transformer architecture framework. The pre-trained model can also be generalized to a diverse set of supervised learning tasks. We pre-trained CodonBERT using 10 million mRNA coding sequences spanning an evolutionarily diverse set of organisms. Next, we used it to perform several mRNA prediction tasks, including protein expression and mRNA degradation prediction. As we show, both the pre-trained and the fine-tuned version of the models can learn new biology and improve on current state-of-the-art methods for mRNA vaccine design.

To assess generalization of our CodonBERT model, we collected a novel hemagglutinin flu vaccine dataset. Different mRNA candidates that encode the influenza hemagglutinin antigen (i.e., with fixed untranslated regions and a variable coding region) were designed, synthesized, and transfected into cells. The protein expression levels corresponding to these mRNA sequences were measured and used as labels for a supervised learning task. CodonBERT leads to better performance than existing methods.

## 2 Results

We developed a large language model, CodonBERT, for mRNA analysis and prediction tasks. CodonBERT was pre-trained using 10 million mRNA sequences derived from mammals, bacteria, and human viruses. All sequences were hierarchically labeled using 13 categories as shown in Figure 1(a). CodonBERT takes the coding region as input using codons as tokens, and outputs an embedding that provides contextual codon representations. The embeddings provided by CodonBERT can be combined with additional trainable layers to perform various downstream regression and prediction tasks, including the prediction of protein expression and mRNA degradation.

**Figure 1:**
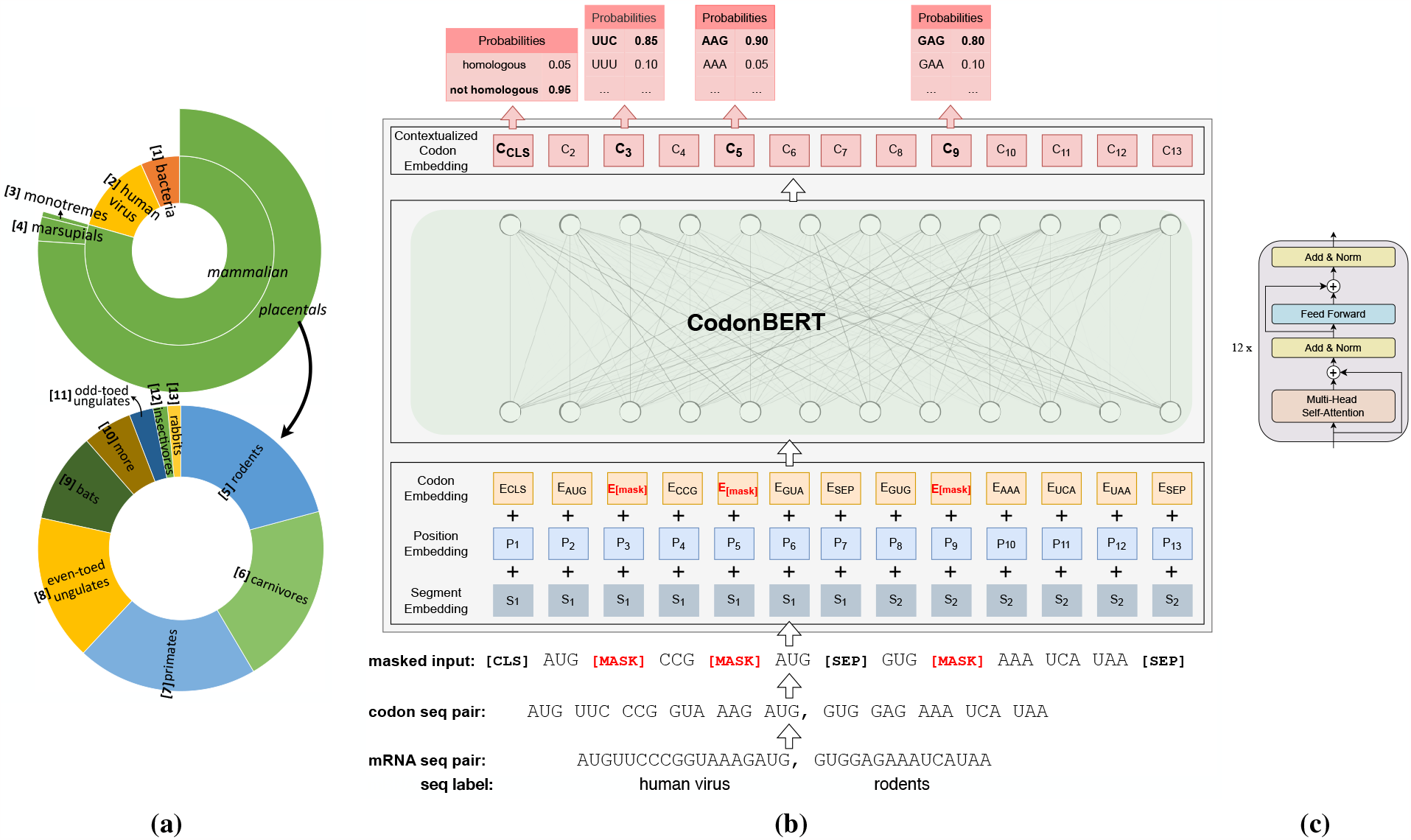
(a) Hierarchically classified mRNA sequences for pre-training. All the 13 leaf-level classes are numbered. The angle of each segment is proportional to the number of sequences belonging to this group. (b) Model architecture and training scheme deployed for two tasks of CodonBERT. (c) A stack of 12 transformer blocks employed in CodonBERT model.

A schematic representation of CodonBERT’s architecture is provided in Figure 1(b). We pre-trained CodonBERT with two tasks: masked language model (MLM) learning and homologous sequences prediction (HSP). The MLM task learns the codon representation, interactions between codons, and relationships between codons and sequences. The HSP task aims to directly model the sequence representation and understand evolutionary relationships between mRNA sequences. In short, a pair of mRNA sequences, which is randomly sampled from either the same or different categories, is codon-tokenized, concatenated, and randomly masked. The masked inputs are further encoded with codon, position, and segment-based embeddings and fed into a stack of transformer layers using a multi-head attention network with residual connections. CodonBERT is self-supervised and relies on masked token prediction and homologous sequence prediction for optimizing parameters (Methods).

### Pre-trained Representation Model

To assess our pre-trained CodonBERT model, we built a held-out dataset by randomly leaving out 1% of mRNA sequences for each category and trained the model with the remaining sequences. As illustrated in Figure S1, during the pre-training phase, the model performance on two tasks (MLM and HSP) is substantially improved on the losses and accuracies of the model prediction on the training and evaluation sets. For example, the entropy loss of the MLM task (*ℒ*_MLM_) decreased to 2.85 on the evaluation set, which means that the model was able to narrow down the choice from 64 (uniform distribution) to only 8 codons for each masked position (Methods).

### CodonBERT learns, on its own, the genetic code and evolutionary homology

In addition to the quantitative evaluation of model predictions, e.g., loss and accuracy, we also performed several qualitative analyses of the embeddings provided by CodonBERT. To decipher what kind of biological information has been learned by the model and encoded in the representation, we randomly sampled 500 sequences for each category from the held-out dataset and extracted high-dimensional codon and sequence embeddings from CodonBERT. These were projected onto a 2-dimensional space by UMAP [37].

In Figure 2(a–b), each dot represents a codon and is annotated with different colors based on its type of codon and amino acid (AA). Codons that encode the same amino acid, i.e., synonymous codons, are spatially close to each other in Figure 2(b), which implies that CodonBERT learns the genetic code from the large-scale training set. For example, the amino acid Valine, whose one letter code is V, can be encoded by four codons: {GUA, GUU, GUG, GUC}. Figure 2(b) illustrates four separate gray clusters for four possible codons, and four clusters are close to each other. We applied the k-nearest neighbors algorithm (kNN) on the output codon embeddings directly. 500 embeddings were sampled for each codon. 99.4% of the 500-nearest neighbors are the same codons. For the remaining misclassified codons, 87.7% are stop codons and are classified to other stop codons.

**Figure 2:**
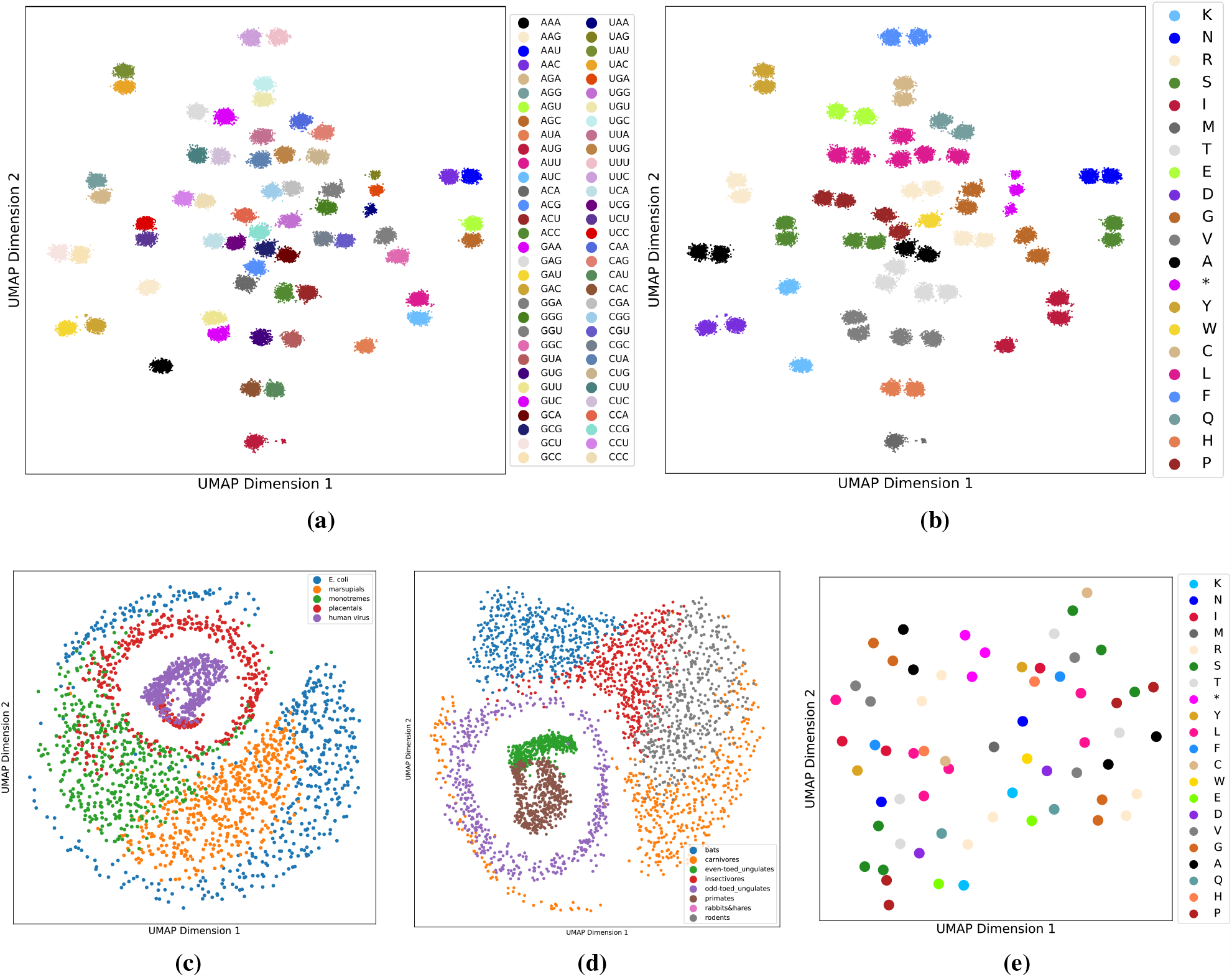
Genetic code and evolutionary homology information learned by pre-trained, unsupervised CodonBERT model. High-dimensional embeddings were projected into 2-dimensional space using UMAP [37]. (a–b) Projected codon embeddings from the pre-trained CodonBERT model. Each point represents a codon with different contexts, and its color corresponds to the type of codon (a) or amino acid accordingly (b). (c–d) Projected sequence embedding from the pre-trained CodonBERT model. Each point is a mRNA sequence, and its color represents the sequence label. (e) Projected codon embedding from the pre-trained Codon2vec model. Each point shows a codon, and its color is the corresponding amino acid.

Codon2vec,a Word2vec [38] model trained on the collected mRNA sequences, can also produce codon representations (Methods). However, when compared to the codon representation generated by CodonBERT, the embedding of each codon from Codon2vec is fixed regardless of the context surrounding the codon. This results in clusters that are often less accurate than the projection of codon representation from CodonBERT (Figure 2e).

In addition to codon representation, CodonBERT also optimizes for sequence identification. 2D projections of the sequence embeddings of the held-out dataset is presented in Figure 2(c–d). Figure 2(c) illustrates clusters of five high-level sequence categories: (*E. coli*, human virus, and three subgroups of mammals). Homologous sequences are clustered together with clear boundaries between the homology classes. As the largest and most developed group within mammals, placentals can be further split into eight specific categories (without the “more” category) in Figure 1(a). As presented in Figure 2(d), clustering of the embeddings corresponding to these eight subgroups is compact and well-separated.

### Evaluating CodonBERT and comparison to prior methods on supervised learning tasks

CodonBERT can be extended to perform supervised learning for specific mRNA prediction tasks. To evaluate the use of our LLM for downstream tasks and to compare it to prior methods, we collected several mRNA prediction datasets. Table 1 presents the datasets and the mRNA properties. As can be seen, these included a diverse set of downstream tasks related to mRNA translation, stability, and regulation. In addition, these datasets represent a range of molecules, including newly published data sets for recombinant protein, bio-computing, and SARS-CoV-2 vaccine design. Finally, we generated a new dataset to test CodonBERT in the context of mRNAs encoding the influenza hemagglutinin antigen for Flu vaccines.

**Table 1:** The collection of the datasets with their corresponding mRNA source and property used for method evaluation. Each dataset is split into training, validation, and test with 0.7, 0.15, 0.15 ratio. All the methods were optimized on the same data split.

| Data Set | Target | category | # mRNAs | seq length |
| --- | --- | --- | --- | --- |
| MLOS Flu Vaccines (Sanofi-Aventis) | Expression | Regression | 543 | 1698 – 1704 |
| mRFP Expression [23] | Expression | Regression | 1459 | 678 – 678 |
| Fungal expression [39] | Expression | Regression | 7056 | 150 – 3000 |
| *E. coli* proteins [40] | Expression | Classification | 6348 | 171 – 3000 |
| Tc-Riboswitches [26] | Switching factor | Regression | 355 | 67 – 73 |
| mRNA stability [41] | Stability | Regression | 41123 | 30 – 1497 |
| SARS-CoV-2 Vaccine Degradation [42] | Degradation | Regression | 2400 | 107 – 107 |

The **mRFP Expression** dataset [23] profiles protein production levels for several gene variants in *E. coli*. The **Fungal expression** dataset [39, 43] includes coding sequences longer than 150 bp from a wide range of fungal genomes. The ***E. coli* proteins** dataset [40] comprises experimental data for protein expression in *E. coli*, which are labeled as low, medium, or high expression (2308, 2067, and 1973 proteins, respectively). The **mRNA stability** dataset [41] includes thousands of mRNA stability profiles obtained from human, mouse, frog, and fish. The **Tc-Riboswitches** dataset [26] consists of a set of tetracycline (Tc) riboswitch dimers sequences upstream of a GFP protein. The measured variable in this data set is the switching factor which refers to the differential effect of the riboswitch in the presence or absence of Tc. The **SARS-CoV-2 Vaccine Degradation** dataset [13] encompasses a set of mRNA sequences that have been tuned for their structural features, stability, and translation efficiency. The benchmarking dataset also included a new dataset generated by Sanofi encoding the hemagglutinin antigen for flu vaccines. Briefly, mRNA sequences, encoding the Influenza H3N2 A/Tasmania/503/2020 hemagglutinin protein, were tested for protein expression level in HeLa cells (Methods).

To compare CodonBERT’s performance on these tasks, we have also applied several other state-of-the-art methods that have been previously used for mRNA property prediction with different model complexities, including TF-IDF [44], TextCNN [45], Codon2vec, RNABERT [34], and RNA-FM [35]. Table 2 presents the performance of CodonBERT and the other six methods on these downstream tasks. For each task, the first three rows are nucleotide-based methods (plain TextCNN, RNABERT, and RNA-FM), while the rest are codon-based methods (TF-IDF, plain TextCNN, Codon2vec, and CodonBERT). Table S1 provides complimentary RMSE values for these comparisons.

**Table 2:** Comparison of CodonBERT to prior methods on seven downstream tasks. For regression tasks, the corresponding Spearman’s rank correlation values are listed. For the classification task (*E. coli* proteins data set), classification accuracy is calculated. The best values of correlation and accuracy for each task are in bold. The corresponding RMSE loss and cross entropy loss is listed in Table S1.

|  | Model | Flu<br>Vaccines | mRFP<br>Expression | Fungal<br>Expression | *E. coli*<br>Proteins | mRNA<br>Stability | Tc-Riboswitch | SARS-CoV-2 Vaccine<br>Degradation |
| --- | --- | --- | --- | --- | --- | --- | --- | --- |
| Nucleotide<br>-based | plain TextCNN | 0.72 | 0.62 | 0.53 | 0.39 | 0.01 | 0.41 | 0.55 |
|  | RNABERT+TextCNN | 0.65 | 0.40 | 0.41 | 0.39 | 0.16 | 0.47 | 0.64 |
|  | RNA-FM+TextCNN | 0.71 | 0.80 | 0.59 | 0.43 | 0.34 | **0.58** | 0.74 |
| Codon<br>-based | TF-IDF | 0.68 | 0.57 | 0.68 | 0.44 | **0.54** | 0.49 | 0.69 |
|  | plain TextCNN | 0.71 | 0.78 | 0.76 | 0.36 | 0.26 | 0.43 | **0.80** |
|  | Codon2vec+TextCNN | 0.72 | 0.77 | 0.61 | 0.43 | 0.33 | 0.56 | 0.70 |
|  | CodonBERT | **0.78** | **0.88** | **0.89** | **0.57** | 0.35 | 0.48 | 0.78 |

Figure S3 ranks different nucleotide- and codon-based methods based on their performance on downstream tasks pertaining to the mRNA-related property prediction. Orange and blue bars represent codon- and nucleotide-based methods, respectively, and the shades correspond to model complexity. Overall, we see that codon-based methods outperform nucleotide-based methods on most tasks. This is in part due to the critical role of codons on the protein expression. Moreover, the codon-based variant of TextCNN outperforms the original nucleotide implementation on most tasks.

As for the detailed comparison, we observe that CodonBERT performed best on four of the seven tasks and second best (in most cases with very small difference) on two of the remaining three tasks. Plain codon-based TextCNN produced the best results for SARS-CoV-2 vaccine degradation, while it performed poorly on other prediction tasks including Riboswitches, Flu vaccines, and *E. coli*. The other two methods that were best performing for one of the datasets, e.g., TF-IDF and RNA-FM, did not perform well on the other tasks.

Both secondary structure and codon usage play critical roles in mRNA vaccine expression [22, 25]. Stable secondary structure increases mRNA stability in solution [25], and optimal codons improve cellular mRNA stability [22]. Therefore, SARS-CoV-2 vaccine degradation and Tc-riboswitch datasets are strongly affected by local and global secondary structure patterns encoded on top of RNA sequences. Although CodonBERT is a codon-based model, it outperforms RNABERT and RNA-FM, which were demonstrated to capture rich structural information from large-scale non-coding RNAs. This may indicate that CodonBERT also learns co-evolutionary information and structural properties from millions of mRNA sequences.

Nucleotide embeddings learned from non-coding RNAs, e.g., RNA-FM_TextCNN_, leads to significantly better results than plain nucleotide-based TextCNN on most tasks. This may indicate that structural information, even from non-coding RNA sequences, is beneficial to solving mRNA translation and stability problems.

Although both RNABERT and RNA-FM are pre-trained BERT models from non-coding RNA sequences, their performance differs. This may be attributed to the training data size and model capacity of RNA-FM which is significantly larger than RNABERT.

## 3 Methods

### Assembly of mRNA Sequences for Pre-training

We collected mRNA sequences across diverse organisms for pre-training from NCBI [46]. The datasets included *mammalian* reference sequences (https://www.ncbi.nlm.nih.gov/datasets/taxonomy/40674/), *bacteria* (*Escherichia coli*) reference sequences (https://www.ncbi.nlm.nih.gov/datasets/taxonomy/562/), and *homo sapiens virus* complete nucleotides (https://www.ncbi.nlm.nih.gov/labs/virus/vssi/#/). Each sequence is with a label representing its taxonomic group.

We pre-processed all the sequences and filtered out some invalid and replicate ones by requiring the mRNA sequences with the sequence length multiples of 3, starting with the start codon (“AUG”) and ending with stop codons (“UAA”, “UAG”, or “UGA”), and only including nucleotides from the set {A, U, G, C, N} (replacing T with U). After pre-processing, 10 million mRNA sequences were valid.

To build sequence pairs for the homologous sequence prediction task, 50% of the sequence pairs consisted of two sequences belonging to one of the 13 categories. The remaining 50% of sequence pairs included two sequences that were randomly sampled from two different categories.

### Model architecture

A codon is composed of three adjacent nucleotides. There are five different options for each of these three positions {A, U, G, C, N} leading to a total of 5^3^ (125) possible combinations. Additionally, five special tokens are added to the vocabulary: classifier token [CLS], separator token [SEP], unknown token [UNK], padding token [PAD], and masking token [MASK]. Thus, in total, there are 130 tokens in the vocabulary of CodonBERT.

As shown in Figure 1(b), CodonBERT takes a sequence pair as input and concatenates them using a separator token ([SEP]). It then adds a classifier token ([CLS]) and a separator token ([SEP]) at the beginning and end of the combined sequence, respectively. CodonBERT constructs the input embedding by concatenating codon, position, and segment embeddings. Absolute positions are utilized with values initialized from 1 to *n*_1_ + *n*_2_ +3 along the concatenated sequence, where *n*_1_ and *n*_2_ are the codon-wise length of two sequences plus three specially added tokens ([CLS] and [SPE]). The segment value is either 1 or 2 to distinguish two sequences. These three types of embedding matrices are learned across 10 millions of mRNA sequences.

The combined input embedding is fed into the CodonBERT model, which consists of a stack of 12 layers of bidirectional transformer encoders [47] as shown in Figure 1(c). Each transformer layer processes its input using 12 self-attention heads, and outputs a representation for each position with hidden size 768. In each layer, the multi-head self-attention mechanism captures the contextual information of the input sequence by considering all the other codons in the sequence. A key benefit of self-attention mechanism is the connection learned between all pairs of positions in an input sequence using parallel computation which enables CodonBERT to model not only short-range but also long-range interactions, which impact translation efficiency and stability [48]. Next a feed-forward neural network is added to apply a non-linear transformation to the output hidden representation from the self-attention network. A residual connection is employed around each of the multi-head attention and feed-forward networks. After processing the input sequence with a stack of transformer encoders, CodonBERT produces the final contextualized codon representations, which is followed by a classification layer to produce probability distribution over the vocabulary during pre-training.

### Pre-training CodonBERT

Model architecture of CodonBERT and the training for the two tasks are illustrated in Figure 1(b). Prior to being fed into the model, the input mRNA sequence is first tokenized into a list of codons. Next, a fraction of input codons (15%) are randomly selected and replaced by the masking token ([MASK]). The self-training loop optimizes CodonBERT to predict the masked codons based on the remaining ones taking into account interactions between the missing and unmasked codons. A probability distribution over 64 possible codons is produced by CodonBERT for the masked positions. The average cross entropy loss *ℒ*_MLM_ over the masked positions *M* is calculated by the optimization function:

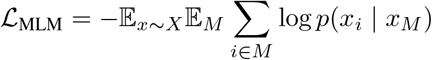

Where *X* represents a batch of sequences, *x* is one sequence and *x*_*i*_ is the original codon for the position *i. x*_*M*_ is the masked input with a set of positions *M* masked. *p*(*x*_*i*_ | *x*_*M*_) indicates the output probability of the real codon *x*_*i*_ given all the remaining codons in the masked sequence *x*_*M*_.

For the HSP task, the output embedding of the classifier token ([CLS]) is used for prediction about whether these two sequences belong to the same class (binary classification). The average cross entropy loss *ℒ*_HSP_ is computed as:

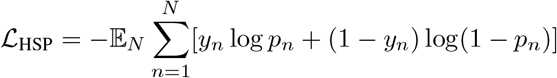

Where *N* represents the number of sequence pairs. *y*_*n*_ is the expected value, which is 1 when two sequences are homologous and 0 when they are not. *p*_*n*_ indicates the predicted probability of two sequences belonging to the same category. The total loss is the sum of the losses from both tasks (*ℒ*_MLM_ + *ℒ*_HSP_).

We used a batch size 128 with sequence length limit 1024, and trained the model around 7 epochs in two weeks. Since the input of CodonBERT are sequence pairs, the length of each sequence is limited up to 512 codons, therefore, the length of the combined sequence is less than 1024. Sequences exceeding the length limitation are split into fragments no longer than 512. Pre-training CodonBERT using 10 million mRNA sequences on 4 A10G GPUs with 96 GB GPU memory and 192 GB memory took roughly two weeks.

CodonBERT was also applied to a wide range of downstream tasks. For this we can use either a single or a pair of sequences as input (Figure 1(b) and Figure 3(c)). To perform supervised analysis, the output embedding is followed by an output layer which is trained for the specific task (protein expression level prediction, mRNA stability, etc.).

**Figure 3:**
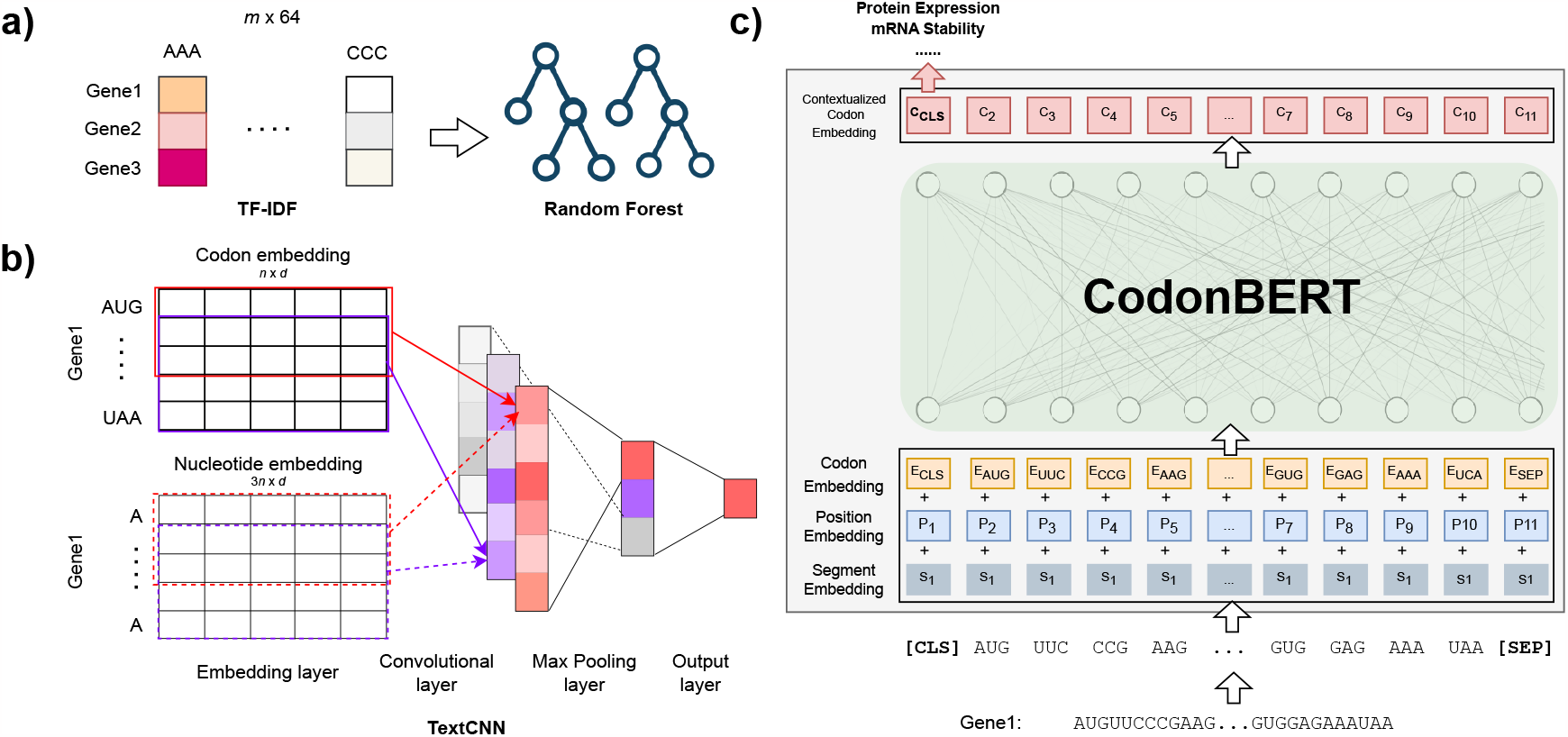
Comparison to prior methods (TF-IDF, Codon2vec, RNABERT and RNA-FM) and fine-tuning CodonBERT on downstream data sets. a) Given an input corpus with *m* mRNA sequences, TF-IDF is used to construct a feature matrix followed by a random forest regression model. b) Use a TextCNN model to learn task-specific nucleotide or codon representations. The model is able to fine-tune pre-trained representations by initializing the embedding layers with stacked codon or nucleotide embeddings extracted from pre-trained language models (Codon2vec, RNABERT, and RNA-FM). *n* is the number of codons in the input sequence and *d* is the dimension of the token embedding. As baseline, plain TextCNN initializes the embedding layer with a standard normal distribution. c) Fine-tune the pre-trained CodonBERT model on a given downstream task directly by keeping all the parameters trainable.

### Pre-training Codon2vec

Word2vec is a popular neural network-based model which is used to learn distributed representations of words in a corpus [38]. Like the large language model mentioned above, Word2vec also learns token representation from a large-scale text corpus, and the embedding from both methods can be utilized as input features for downstream tasks. Unlike LLMs it only produces a single assignment for each token, which usually correlates with the max or most frequent context around the token in the corpus.

An existing work has applied Word2vec to the fungal genomes, studied codon usage bias, and built a predictive model for gene expression [39]. However, there is no application on large-scale mRNA sequence dataset. Therefore, for comparison, we trained our own Codon2vec model on the collected mRNA sequences. Using the Gensim library [49] the Codon2vec model opted for a skip-gram architecture, accompanied by a window size of five and a minimum count threshold of 10 codons. The model was trained using hierarchical SoftMax and negative sampling methodologies.

The input sequences for pre-training Codon2vec were processed through a filtration system that selected sequences containing fewer than 1000 nucleotides. This filtration stage was necessitated by the constrained model capacity of the Word2vec neural network. Upon completion of this process, we retrieved a total of 2,301,807 sequences, which were subsequently subjected to tokenization into k-mers. These k-mers serve as representations of all corresponding codons.

### *In vitro* transcription, cell culture, and transfections

As shown in Figure S4, mRNA sequences were designed using to encode the Influenza H3N2 A/Tasmania/503/2020 hemagglutinin protein. Sequences corresponding to these candidates were synthesized as gene fragments and PCR amplified to generate template DNA for High Throughput *in vitro* transcription reaction containing N1-Methylpseudouridine. The resulting purified precursor mRNA was reacted further via enzymatic addition of a 5*′* cap structure (Cap 1) and a 3*′* poly(A) tail of approximately 200 nucleotides in length as determined by capillary electrophoresis.

HeLa cells were used the evaluate the expression of the protein encoded by different mRNA sequences. Cells were cultured and maintained in MEM (Corning) containing 10% (v/v) heat inactivated FBS (Gibco). To evaluate the expression of candidate mRNAs, HeLa were transiently transfected with mRNAs complexed with Lipofectamine MessengerMax (Thermo Scientific). Unknown mRNAs were thawed, diluted in Opti-MEM, combined with Lipofectamine for 10 minutes, and were then further diluted in Opti-MEM. To prepare the cells for reverse transfection, HeLa were collected from culture flasks using TrypLE and were diluted in complete growth medium such that each well will be seeded with 2E4 live cells. Complexed mRNAs (20 ng/well) were added to triplicate wells of a 96-well poly-D-lysine PhenoPlate (PerkinElmer) and were combined with 2E4 HeLa cells. Plates were rested at RT briefly before incubation in a tissue culture incubator for 20 hours + 30 minutes. At the endpoint, cells were lysed in RIPA (Thermo Scientific) supplemented with Omnicleave (Lucigen) and HALT protease inhibitor (Thermo Scientific). The hemagglutinin expression in cell lysates was determined using a quantitative sandwich ELISA, and the expression level of unknown mRNAs was normalized to the value from a known benchmark mRNA sequence.

### Comparisons to other prior methods

We compared CodonBERT to several prior methods that have been used to model and analyze RNA sequences.

- Term Frequency-Inverse Document Frequency (TF-IDF) [44] is a numerical statistic that is commonly used as a weighting scheme in information retrieval and natural language processing. In the context of mRNA sequences, TF-IDF is applied to measure the significance of each codon in a sequence. High TF-IDF value of a codon indicates that the codon is important in a particular mRNA sequence, and rare across all mRNA sequences in the corpus.
- Convolutional Neural Network (CNN) was first proposed and has been commonly used in image recognition and was later also applied for text analysis TextCNN [45]. TextCNN consists of multiple types of layers including an embedding layer, convolutional layer, pooling layer and fully connected layer shown in Figure 3(b). Each row of the embedding layer represents a token.
- RNABERT [34] and RNA-FM [35] are RNA large language models. However, they are pre-trained on non-coding RNAs to learn and encode structural and functional properties in the output nucleotide embedding.

## 4 Discussion

To enable the analysis and prediction of mRNA properties, we utilized 10 million mRNA coding sequences (CDS) from several species to train a large language model (CodonBERT), and to establish a foundational model. The model optimizes two self-supervised tasks: codon completion and homology detection. Like other unsupervised LLMs, we expected that such a foundational model will learn to capture aspects of natural selection that favors mRNA sequences with high expression and stable structure. Analysis of the resulting model indicates that it indeed learns several relevant biological properties for codons and sequences.

Projection of codon embedding obtained from CodonBERT produces distinct clusters that adhere to the amino acid types. In-depth analysis of CodonBERT representation of a set of genes from different organisms revealed that CodonBERT autonomously learns the genetic code and principles of evolutionary homology. The projection of the clusters leads to a spiral shape (Figure 2(c–d)) which separates organisms as well as genes based on their functions. This may indicate that CodonBERT not only learns the sequence of evolutionary occurrence but can learn a pseudo evolutionary tree as part of its embeddings.

Further, the sequence representations and clusters obtained from CodonBERT align with our understanding of genetic evolution. While the genetic information across these species is different, indicating diversity, there is also evidence of parallel evolution, where genes have evolved in a similar trajectory amongst these species. This parallel evolution maintains their shared functions and ensures their genetic distinctness simultaneously.

We also utilized CodonBERT to perform several supervised prediction tasks for mRNA properties. These include datasets testing for recombinant protein expression, mRNA degradation, mRNA stability, and more. Our results indicate that CodonBERT is the top performing method overall and ranks first or second in performance for six of the seven tasks. All other methods we compared to performed poorly on all, or some of the tasks. Thus, for a new task, the use of CodonBERT is likely to lead to either the best or close to best results. CodonBERT’s success in the more structurally related tasks (including mRNA stability and the TC riboswitches datasets) indicates that it can learn co-evolutionary and structural concepts using large-scale mRNA sequences.

For the vaccine-related downstream tasks, CodonBERT generally exhibited robust performance. It was 10% better than the second-best method for the new hemagglutinin flu vaccine expression dataset and very close (2% difference) to the top performing model for the SARS-CoV-2 Vaccine degradation dataset. Although fungal sequences are not included in pre-training sequences, CodonBERT model shows its generalization to a downstream fungal dataset. RNABERT and RNABERT are also pre-trained RNA large language models; however, they are nucleotide-based and trained on non-coding RNAs. Since they do not explicitly capture codon usage, these methods are inferior to CodonBERT on the protein expression-related tasks.

The one exception in terms of performance was observed for the mRNA stability task. Stability is known to be structure-dependent, and stable structures such as stem-loops or hairpin structures can impede degradation enzymes, protecting the mRNA from rapid decay. A possible reason for the reduction in performance for this dataset is that structural properties are highly dependent on nucleotides whereas CodonBERT is a codon-based model. One possible solution for this is a model that combines codon and nucleotide representation. Similarly, mRNA modification events including capping at the 5*′* end and polyadenylation at the 3*′* end in eukaryotes are not currently encoded in our model but can also impact mRNA stability.

While the tasks we have focused on are supervised in nature, CodonBERT, as an LLM, can be utilized for generative purposes. Specifically, we envision using this model for codon optimization of various heterologous proteins for vaccines. Other generative tasks can include sampling mRNA for synthetic biology use cases (e.g., creation of optimized mRNA constructs for genome editing).

To conclude, our findings suggest that CodonBERT could serve as a versatile and foundational model for the development of new mRNA-based vaccines and the engineering and recombinant production of industrial and therapeutic proteins.

## References

[1] N. Pardi, M. J. Hogan, F. W. Porter, and D. Weissman, “mRNA vaccines — a new era in vaccinology,” Nature Reviews Drug Discovery, vol. 17, no. 4, pp. 261–279, 2018.

[2] N. Pardi, M. J. Hogan, and D. Weissman, “Recent advances in mRNA vaccine technology,” Current opinion in immunology, vol. 65, pp. 14–20, 2020.

[3] N. A. Jackson, K. E. Kester, D. Casimiro, S. Gurunathan, and F. DeRosa, “The promise of mRNA vaccines: a biotech and industrial perspective,” npj Vaccines, vol. 5, no. 1, p. 11, 2020.

[4] C. Zhang, G. Maruggi, H. Shan, and J. Li, “Advances in mRNA vaccines for infectious diseases,” Frontiers in immunology, p. 594, 2019.

[5] L. A. Jackson, E. J. Anderson, N. G. Rouphael, P. C. Roberts, M. Makhene, R. N. Coler, M. P. McCullough, J. D. Chappell, M. R. Denison, L. J. Stevens, et al., “An mRNA vaccine against SARS-CoV-2—preliminary report,” New England journal of medicine, vol. 383, no. 20, pp. 1920–1931, 2020.

[6] E. H. Pilkington, E. J. Suys, N. L. Trevaskis, A. K. Wheatley, D. Zukancic, A. Algarni, H. Al-Wassiti, T. P. Davis, C. W. Pouton, S. J. Kent, et al., “From influenza to COVID-19: Lipid nanoparticle mRNA vaccines at the frontiers of infectious diseases,” Acta biomaterialia, vol. 131, pp. 16–40, 2021.

[7] N. Pardi, M. J. Hogan, R. S. Pelc, H. Muramatsu, H. Andersen, C. R. DeMaso, K. A. Dowd, L. L. Sutherland, R. M. Scearce, R. Parks, et al., “Zika virus protection by a single low-dose nucleoside-modified mrna vaccination,” Nature, vol. 543, no. 7644, pp. 248–251, 2017.

[8] G. Maruggi, E. Chiarot, C. Giovani, S. Buccato, S. Bonacci, E. Frigimelica, I. Margarit, A. Geall, G. Bensi, and D. Maione, “Immunogenicity and protective efficacy induced by self-amplifying mRNA vaccines encoding bacterial antigens,” Vaccine, vol. 35, no. 2, pp. 361–368, 2017.

[9] L. Miao, Y. Zhang, and L. Huang, “mRNA vaccine for cancer immunotherapy,” Molecular Cancer, vol. 20, no. 1, pp. 1–23, 2021.

[10] C. L. Lorentzen, J. B. Haanen, Ö. Met, and I. M. Svane, “Clinical advances and ongoing trials on mRNA vaccines for cancer treatment,” The Lancet Oncology, vol. 23, no. 10, pp. e450–e458, 2022.

[11] T. Schlake, A. Thess, M. Fotin-Mleczek, and K.-J. Kallen, “Developing mRNA-vaccine technologies,” RNA Biology, vol. 9, no. 11, pp. 1319–1330, 2012. PMID: 23064118.

[12] H. I. Ahmad, A. Jabbar, N. Mushtaq, Z. Javed, M. U. Hayyat, J. Bashir, I. Naseeb, Z. U. Abideen, N. Ahmad, and J. Chen, “Immune tolerance vs. immune resistance: The interaction between host and pathogens in infectious diseases,” Frontiers in Veterinary Science, vol. 9, p. 827407, 2022.

[13] K. Leppek, G. W. Byeon, W. Kladwang, H. K. Wayment-Steele, C. H. Kerr, A. F. Xu, D. S. Kim, V. V. Topkar, C. Choe, D. Rothschild, G. C. Tiu, R. Wellington-Oguri, K. Fujii, E. Sharma, A. M. Watkins, J. J. Nicol, J. Romano, B. Tunguz, F. Diaz, H. Cai, P. Guo, J. Wu, F. Meng, S. Shi, E. Participants, P. R. Dormitzer, A. Solórzano, M. Barna, and R. Das, “Combinatorial optimization of mRNA structure, stability, and translation for RNA-based therapeutics,” Nature Communications, vol. 13, no. 1, p. 1536, 2022.

[14] V. P. Mauro and S. A. Chappell, “A critical analysis of codon optimization in human therapeutics,” Trends in Molecular Medicine, vol. 20, no. 11, pp. 604–613, 2014.

[15] A. B. Al-Hawash, X. Zhang, and F. Ma, “Strategies of codon optimization for high-level heterologous protein expression in microbial expression systems,” Gene Reports, vol. 9, pp. 46–53, 2017.

[16] G. R. Webster, A. Y.-H. Teh, and J. K.-C. Ma, “Synthetic gene design—the rationale for codon optimization and implications for molecular pharming in plants,” Biotechnology and Bioengineering, vol. 114, no. 3, pp. 492–502, 2017.

[17] V. P. Mauro, “Codon optimization in the production of recombinant biotherapeutics: Potential risks and considerations,” BioDrugs, vol. 32, no. 1, pp. 69–81, 2018.

[18] A. H. Parret, H. Besir, and R. Meijers, “Critical reflections on synthetic gene design for recombinant protein expression,” Current Opinion in Structural Biology, vol. 38, pp. 155–162, 2016.

[19] M. Schmidt, K. Hamacher, F. Reinhardt, T. S. Lotz, F. Groher, B. Suess, and S. Jager, “SICOR: Subgraph isomorphism comparison of rna secondary structures,” IEEE/ACM Transactions on Computational Biology and Bioinformatics, vol. 17, no. 6, pp. 2189–2195, 2020.

[20] F. Groher, C. Bofill-Bosch, C. Schneider, J. Braun, S. Jager, K. Geißler, K. Hamacher, and B. Suess, “Riboswitching with ciprofloxacin—development and characterization of a novel RNA regulator,” Nucleic Acids Research, vol. 46, pp. 2121–2132, 01 2018.

[21] V. Agarwal and J. Shendure, “Predicting mRNA abundance directly from genomic sequence using deep convolutional neural networks,” Cell reports, vol. 31, no. 7, 2020.

[22] V. Agarwal and D. R. Kelley, “The genetic and biochemical determinants of mRNA degradation rates in mammals,” Genome Biology, vol. 23, no. 1, p. 245, 2022.

[23] T. Nieuwkoop, B. R. Terlouw, K. G. Stevens, R. Scheltema, D. de Ridder, J. van der Oost, and N. Claassens, “Revealing determinants of translation efficiency via whole-gene codon randomization and machine learning,” Nucleic Acids Research, vol. 51, pp. 2363–2376, 01 2023.

[24] H. Zhang, L. Zhang, A. Lin, C. Xu, Z. Li, K. Liu, B. Liu, X. Ma, F. Zhao, H. Jiang, C. Chen, H. Shen, H. Li, D. H. Mathews, Y. Zhang, and L. Huang, “Algorithm for optimized mRNA design improves stability and immunogenicity,” Nature, 2023.

[25] D. M. Mauger, B. J. Cabral, V. Presnyak, S. V. Su, D. W. Reid, B. Goodman, K. Link, N. Khatwani, J. Reynders, M. J. Moore, and I. J. McFadyen, “mRNA structure regulates protein expression through changes in functional half-life,” Proceedings of the National Academy of Sciences, vol. 116, no. 48, pp. 24075–24083, 2019.

[26] A.-C. Groher, S. Jager, C. Schneider, F. Groher, K. Hamacher, and B. Suess, “Tuning the performance of synthetic riboswitches using machine learning,” ACS Synthetic Biology, vol. 8, no. 1, pp. 34–44, 2019.

[27] S. Li, H. Zhang, L. Zhang, K. Liu, B. Liu, D. H. Mathews, and L. Huang, “LinearTurboFold: linear-time global prediction of conserved structures for RNA homologs with applications to SARS-CoV-2,” Proceedings of the National Academy of Sciences, vol. 118, no. 52, p. e2116269118, 2021.

[28] M. E. Peters, M. Neumann, M. Iyyer, M. Gardner, C. Clark, K. Lee, and L. Zettlemoyer, “Deep contextualized word representations,” in North American Chapter of the Association for Computational Linguistics, 2018.

[29] A. Radford, K. Narasimhan, T. Salimans, I. Sutskever, et al., “Improving language understanding by generative pre-training,” 2018.

[30] J. Devlin, M.-W. Chang, K. Lee, and K. Toutanova, “Bert: Pre-training of deep bidirectional transformers for language understanding,” arXiv preprint arXiv:1810.04805, 2018.

[31] T. Shen, Z. Hu, Z. Peng, J. Chen, P. Xiong, L. Hong, L. Zheng, Y. Wang, I. King, S. Wang, et al., “E2Efold-3D: End-to-End Deep Learning Method for accurate de novo RNA 3D Structure Prediction,” arXiv preprint arXiv:2207.01586, 2022.

[32] A. Rives, J. Meier, T. Sercu, S. Goyal, Z. Lin, J. Liu, D. Guo, M. Ott, C. L. Zitnick, J. Ma, et al., “Biological structure and function emerge from scaling unsupervised learning to 250 million protein sequences,” Proceedings of the National Academy of Sciences, vol. 118, no. 15, p. e2016239118, 2021.

[33] T. Bepler and B. Berger, “Learning the protein language: Evolution, structure, and function,” Cell systems, vol. 12, no. 6, pp. 654–669, 2021.

[34] M. Akiyama and Y. Sakakibara, “Informative RNA base embedding for RNA structural alignment and clustering by deep representation learning,” NAR genomics and bioinformatics, vol. 4, no. 1, p. qac012, 2022.

[35] J. Chen, Z. Hu, S. Sun, Q. Tan, Y. Wang, Q. Yu, L. Zong, L. Hong, J. Xiao, T. Shen, I. King, and Y. Li, “Interpretable RNA foundation model from unannotated data for highly accurate RNA structure and function predictions,” bioRxiv, 2022.

[36] Y. Ji, Z. Zhou, H. Liu, and R. V. Davuluri, “DNABERT: pre-trained Bidirectional Encoder Representations from Transformers model for DNA-language in genome,” Bioinformatics, vol. 37, pp. 2112–2120, 02 2021.

[37] L. McInnes, J. Healy, and J. Melville, “UMAP: Uniform manifold approximation and projection for dimension reduction,” 2020.

[38] T. Mikolov, K. Chen, G. Corrado, and J. Dean, “Efficient estimation of word representations in vector space,” arXiv preprint arXiv:1301.3781, 2013.

[39] R. Wint, A. Salamov, and I. V. Grigoriev, “Kingdom-Wide Analysis of Fungal Protein-Coding and tRNA Genes Reveals Conserved Patterns of Adaptive Evolution,” Molecular Biology and Evolution, vol. 39, 01 2022. msab372.

[40] Z. Ding, F. Guan, G. Xu, Y. Wang, Y. Yan, W. Zhang, N. Wu, B. Yao, H. Huang, T. Tuller, et al., “MPEPE, a predictive approach to improve protein expression in E. coli based on deep learning,” Computational and Structural Biotechnology Journal, vol. 20, pp. 1142–1153, 2022.

[41] M. Diez, S. G. Medina-Muñoz, L. A. Castellano, G. da Silva Pescador, Q. Wu, and A. A. Bazzini, “iCodon customizes gene expression based on the codon composition,” Scientific Reports, vol. 12, no. 1, pp. 1–16, 2022.

[42] H. K. Wayment-Steele, W. Kladwang, A. M. Watkins, D. S. Kim, B. Tunguz, W. Reade, M. Demkin, J. Romano, R. Wellington-Oguri, J. J. Nicol, et al., “Deep learning models for predicting RNA degradation via dual crowdsourcing,” Nature Machine Intelligence, pp. 1–11, 2022.

[43] I. V. Grigoriev, R. Nikitin, S. Haridas, A. Kuo, R. Ohm, R. Otillar, R. Riley, A. Salamov, X. Zhao, F. Korzeniewski, et al., “Mycocosm portal: gearing up for 1000 fungal genomes,” Nucleic acids research, vol. 42, no. D1, pp. D699–D704, 2014.

[44] A. Rajaraman and J. D. Ullman, Mining of massive datasets. Cambridge University Press, 2011.

[45] Y. Kim, “Convolutional neural networks for sentence classification,” 2014.

[46] D. L. Wheeler, T. Barrett, D. A. Benson, S. H. Bryant, K. Canese, V. Chetvernin, D. M. Church, M. DiCuccio, R. Edgar, S. Federhen, et al., “Database resources of the national center for biotechnology information,” Nucleic acids research, vol. 35, no. suppl_1, pp. D5–D12, 2007.

[47] A. Vaswani, N. Shazeer, N. Parmar, J. Uszkoreit, L. Jones, A. N. Gomez, Ł. Kaiser, and I. Polosukhin, “Attention is all you need,” Advances in neural information processing systems, vol. 30, 2017.

[48] J. G. A. Aw, Y. Shen, A. Wilm, M. Sun, X. N. Lim, K.-L. Boon, S. Tapsin, Y.-S. Chan, C.-P. Tan, A. Y. Sim, et al., “In vivo mapping of eukaryotic RNA interactomes reveals principles of higher-order organization and regulation,” Molecular cell, vol. 62, no. 4, pp. 603–617, 2016.

[49] R. Rehurek and P. Sojka, “Gensim–python framework for vector space modelling,” NLP Centre, Faculty of Informatics, Masaryk University, Brno, Czech Republic, vol. 3, no. 2, 2011.

