## Supplemental Information for "CodonBERT: Large Language Models for mRNA design and optimization"

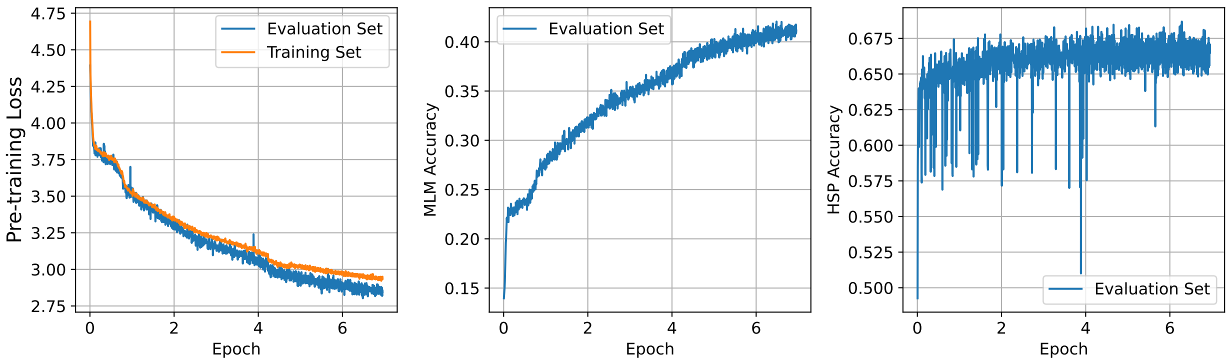


**Figure S1:** The pre-training curve of CodonBERT.

| Model / Dataset | Flu Vaccines | mRFP Expression | Fungal Expression | *E. coli* Proteins | mRNA Stability | Tc-Riboswitch | CoV Vaccine Degradation |
|---|---|---|---|---|---|---|---|
| number of seqs | 538 | 1459 | 7553 | 6348 | 41123 | 355 | 2400 |
| plain TextCNN | **0.26** / 0.72 | 0.35 / 0.62 | 3.35 / 0.53 | 1.09 / 0.39 | 1.01 / 0.01 | 0.53 / 0.41 | 0.017 / 0.55 |
| RNABERT | 0.45 / 0.65 | 0.47 / 0.40 | 3.85 / 0.41 | 1.09 / 0.39 | 0.98 / 0.16 | 0.44 / 0.47 | 0.017 / 0.64 |
| RNA-FM | 0.36 / 0.71 | 0.21 / 0.80 | 3.06 / 0.59 | 1.05 / 0.43 | 0.89 / 0.34 | 0.45 / **0.58** | 0.015 / 0.74 |
| TF-IDF | 0.37 / 0.68 | 0.43 / 0.57 | 2.59 / 0.68 | – / 0.44 | 0.68 / **0.54** | 0.46 / 0.49 | 0.017 / 0.69 |
| plain TextCNN | 0.37 / 0.71 | 0.21 / 0.78 | 1.83 / 0.76 | 1.09 / 0.36 | **0.59** / 0.26 | 0.64 / 0.43 | **0.009 / 0.80** |
| Codon2vec$_{+\text{TextCNN}}$ | 0.30 / 0.72 | 0.28 / 0.77 | 3.04 / 0.61 | 1.06 / 0.43 | 0.91 / 0.33 | **0.43** / 0.56 | 0.016 / 0.70 |
| CodonBERT | 0.28 / **0.78** | **0.11 / 0.88** | **0.64 / 0.89** | **0.92 / 0.57** | 0.94 / 0.35 | 0.44 / 0.48 | 0.012 / 0.78 |

**Table S1:** Results of our CodonBERT model against other benchmarks on the test set of seven downstream tasks. For regression tasks, the RMSE loss and the corresponding Spearman's rank correlation are listed. For the classification task (*E. coli* proteins data set), the cross entropy loss and classification accuracy are calculated. The best values of loss, correlation and accuracy for each task are in bold. Fig. S2 shows scatter plots of predicted protein expression values from CodonBERT and other existing benchmarks against experimental values.

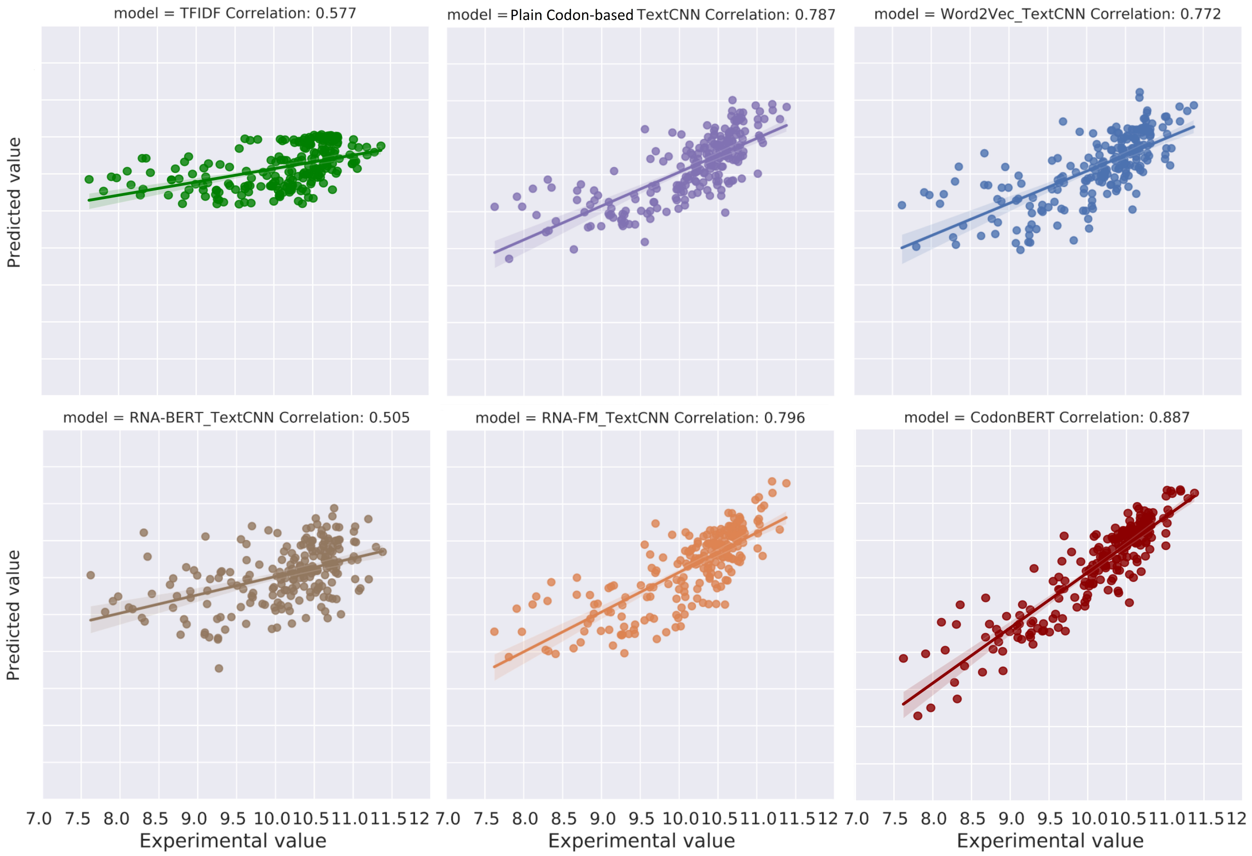


**Figure S2:** Scatter plots of predicted protein expression values from CodonBERT and other existing benchmarks against experimental values.

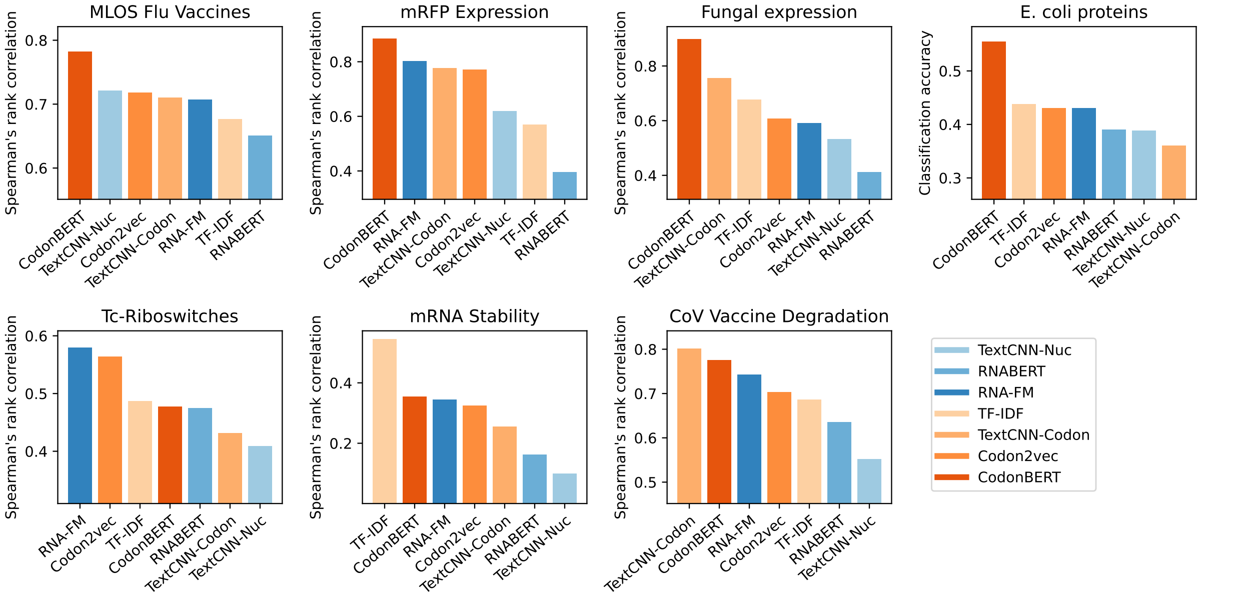


**Figure S3:** Ranked results (Spearman's rank correlation or classification accuracy, higher is better) for CodonBERT model and other benchmarks. Nucleotide-based and codon-based methods are in blue and orange colors, respectively. Shades represent model complexity.

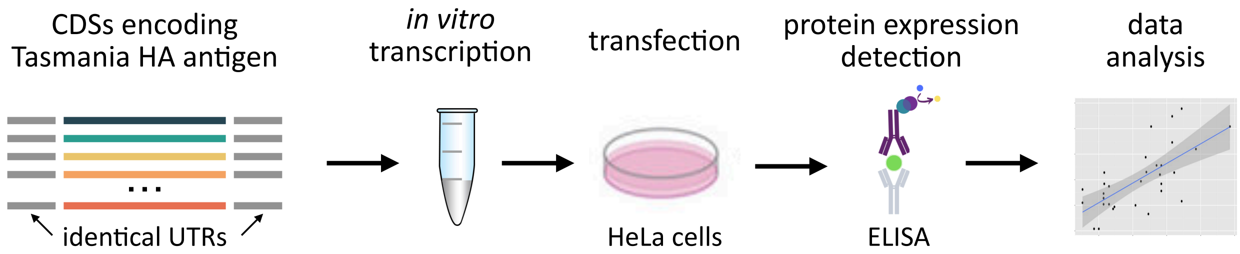


**Figure S4:** Experimental design for testing in-cell protein expression.
